# Pentilludin reduces rat amphetamine and remifentail self-administration with good pharmacologic and toxicologic profiles

**DOI:** 10.1101/2025.07.28.667221

**Authors:** George Uhl, Balaji Kannan, Joungil Choi, Ian Henderson, Bridget Gregory, Jared Solon, Corinne Wells, Edward D Levin

## Abstract

Pentilludin is a novel, potent (690 nM) irreversible inhibitor of actions of the receptor type protein tyrosine phosphatase D (PTPRD). Pentilludin displays no *in vitro* activities in Ames or micronucleus tests, at hERG channels or at targets for currently-licensed drugs. Rats treated with pentilludin doses up to 100 mg/kg/day for two weeks have not been found to display behavioral, hematologic or serum chemistry abnormalities. Treatment with 20 mg/kg sc pentilludin prior to every other M-W-F self-administration session substantially reduces self-administration of amphetamine and more modestly reduces self-administration of remifentanil. Pentilludin provides a novel means for reducing self-administration of psychostimulant and, modestly, opiate drugs in ways that could enhance abstinence in humans.

## Introduction

Pentilludin (NHB1109) is a 7-position substituted cyclopentyl methoxy illudalic acid analog that potently and (likely) irreversibly inhibits actions of the receptor type protein tyrosine phosphatase PTPRD (Fig 1; (1-3). We identified pentilludin after modeling *in silico*, synthesizing and testing > 70 novel candidate PTPRD ligands, many related to our lead compound PTPRD phosphatase inhibitor, 7-BIA (4). Pentilludin displays 690 nM potency in inhibiting PTPRD’s phosphatase, likely irreversibly. It displays some potency at closely related receptor type protein tyrosine phosphatases S and F (PTPRS and PTPRF), and less potency at the receptor type protein tyrosine phosphatase J and the nonreceptor type protein tyrosine phosphatase 1 (PTPRJ and PTPN1/ PTP1B)(2). IC_50_ values are ≥10^−4^ M at all targets of the NIDA-EUROFINS assays that include virtually all targets of currently marketed drugs (1).

**Fig 1.**
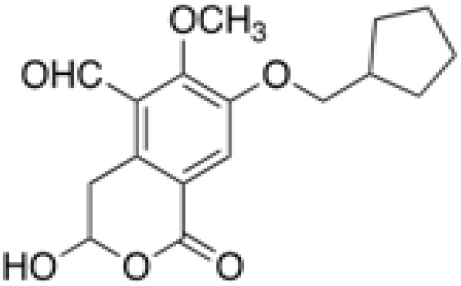
Structure of pentilludin (7-methoxy-cyclopentyl illudalic acid analog; NHB1109 (1)

Several lines of evidence suggest that a drug that reduced PTPRD activity would provide safe and useful actions related to substance use disorders. Heterozygous knockout mice with 50% of wildtype levels of expression display less reward from psychostimulant administration, as assessed in conditioned place preference or self-administration assays (4, 5). These mice are otherwise similar to wildtype littermates in tests of acute nociception (hotplate, tailflick), memory (Morris water maze), fear/anxiety (dark box emergence, thigmotaxis), and motor abilities (screen hang time, locomotion, rotarod) (5). Human genetic results (6) associate common variation in PTPRD with: 1) vulnerability to develop a substance use disorder (polysubstance use (7-9), opioid use disorder (10) and alcohol use disorder (11)); 2) ability to quit smoking (12, 13); 3) ability to quit use of opioids (14); 4) ability to reduce alcohol use (when aided by naltrexone) (15); and 5) levels of expression of PTPRD mRNA in human postmortem cortex (5). PTPRD SNPs display 10^−6^ < p < 10^−7^ association with a specific constellation of rewarding responses to amphetamine administered po in a human laboratory setting (16)(17).

Drugs that sharply reduce PTPRD activity reduce appetite, providing dose-limiting toxicity in mice (1). PTPRD serves as the physiological hypothalamic receptor of orexigenic activities of the fat cell orexigenic hormone asprosin (18). Homozygous PTPRD knockout mice are small and require moistened food placement on the floors of their cages for optimal survival until adulthood (5). While no weight loss was seen with repeated doses of 100 mg/kg/day, mice treated daily with ≥ 150 mg/kg pentilludin po reduce oral intake and lose weight starting on about the 4^th^ day of repeated daily dosing.

Rats are often used for studies that characterize properties of drug candidates and frequently provide data that support use in humans (19). Self-administration of addictive substances by rats provides a highly-developed platform for testing the abilities of potential therapeutics to reduce drug reward. We thus now report a) studies of pentilludin pharmaco- and toxicokinetics that validate its use of rats at doses up to 100-150 mg/kg/d po and b) tests of effects of intermittent pentilludin dosing at 20 mg/kg sc on self-administration of a stimulant, amphetamine, and an opiate agonist, remifentanil. Taken together, these and control/comparison results a) support use of rat models for studies of pentilludin toxicity and efficacy and b) encourage development of pentilludin as a therapeutic in settings where pharmacologic reduction in reward from stimulants, and possibly opiates, is desirable.

## Materials and Methods

### *Pentilludin was synthesized* as described (1)

Pentilludin powder was >97.5% pure following synthesis and when reanalyzed (Certificate of Analysis, GLP) 1 ½ and 2 ½ years after synthesis. Pentilludin was soluble in 6.82% Captisol in 60:40 PEG600/water (the vehicle used for initial toxicology experiments) and in PEG600 (the vehicle used for self-administration and Good Laboratory Practices testing) at concentrations up to 100 mg/ml with gentle heating.

### IACUC approvals

All animal studies were performed under protocols approved by the relevant Institutional Animal Care and Use Committee reviews (Duke University School of Medicine, CARE/Mountain West Research LLC and Bioduro-Sundia).

### *Method for measuring pentilludin* in rat plasma

Proteins in 50 μL plasma collected with K_2_EDTA as anti-coagulant were precipitated by adding 5 μL of acetonitrile with 0.5% v/v ascorbate, 200 μL of acetonitrile containing 20 ng/ml tolbutamide internal standard with 0.5% ascorbate was added, the mixture was vortexed for 1 min, centrifuged @ 4000 rpm for 15 mins and peaks in 8 μL of the supernatant were identified using an AB Sciex Triple Quad 5500 LC-MS/MS (electrospray, negative ions, Analyst 1.6.3 software) and a Waters Acquity Ultra Performance LC System with 50 × 2.1 mm Acuity UPLC Beh C18 1.7 μm column eluted for 2.5 min @ 22^°^C @ 0.5 ml/min with Mobile Phase A (5mM NH_4_OAc in water with 0.05% formic acid) and mobile phase B (acetonitrile with 0.1% formic acid) with changes in % A as follows: 60, 10, 5, and 60. Standards were quantitated at concentrations ranging from 1-2000 ng/mL.

### Methods for assessing pentilludin toxicities

Observational battery: Rats were assessed twice daily using an observational battery under Good Laboratory Practice conditions that assessed evidence for abnormalities related to convulsions, tremors, somasthenia, posture at rest and with movement, breathing, body temperature, weight loss, condition of genitalia, skin and hair as well as quality and quantity of secretions, urine and feces (nonGLP) and eye appearance, lacrimation, muscle tone, salivation, piloerection, fur appearance, body temperature, stool appearance, tonic movements, clonic movements, fasciculations, tremors, posture, gait, arousal, urine and fecal output and rearing.

Blood was collected for assessment of standard hematological and chemistry parameters. Tissues were collected at necropsy for histopathological analyses by Veterinary pathologists of hemotoxylin/ eosin-stained paraffin embedded section from aorta, heart, lung with mainstem bronchi, esophagus, tongue, trachea, sciatic nerve, skeletal muscle, sternal bone with marrow, eye with optic nerve, Harderian gland, brain, pituitary and cervical, thoracic and lumbar spinal cord. These studies were performed at Experimental Pathology Laboratories and Bioduro.

### Method for amphetamine and remifentanil self-administration

Young adult Sprague-Dawley rats (Charles River Laboratories, Raleigh, NC, USA) were singly housed on a reverse 12:12 day night cycle and tested during their active (dark) phase in studies approved by the Duke Institutional Animal Care and Use Committee. Rats had catheters surgically implanted in their jugular veins to provide access for self-administration by IV infusion. Briefly, rats were anesthetized with i.p. injections of ketamine (Fort Dodge Animal Health, Fort Dodge, IA, USA; 0.6 mg/kg) with the addition of Dexdomitor (Pfizer, New York, NY, USA; 0.15 mg/kg) which is a sedative and analgesic agent. Before surgery, ketoprofen (5 mg/kg, s.c.) was administered to reduce post-operative pain and inflammation. A plastic SoloPort was attached intraoperatively to a polyurethane catheter and inserted into a subcutaneous interscapular pocket and sutured to underlying fascia.

The day following surgery, the rats began self-administration sessions with amphetamine (0.1 mg/kg/infusion, iv) or remifentanil (0.3 µg/kg/infusion, iv) as reinforcers. A lever press on the active side lever resulted in the activation of the feedback tone for 0.5 sec and the immediate delivery of one 50-µl infusion of drug in less than 1 sec. Each infusion was followed by a 20-sec timeout in which the house light goes on and cue light goes out. Responses were recorded but not reinforced during this interval. Inactive lever presses were recorded but resulted in no infusion. There were five sessions of training for self-administration before the onset of pentilludin drug treatment sessions. All of the animals had the same number of training sessions to provide all with equal testing history. Each infusion session lasted for one hour. Animals were tested not more than once/day between 9 AM and 4 PM.

At the end of each study catheter patency was confirmed by assessing sedation following methohexital (brevital) administration through the catheter. Only those rats with verified patent catheters were included in statistical analysis. Self-administration results, differences from baseline levels, were assessed for statistical for significance with analysis of variance. To control for baseline differences in self-administration, the self-administration with pentilludin or vehicle treatment was analyzed as a difference from each animal’s self-administration on the last two sessions of baseline training. The self-administration studies were run in repeated cohorts, which was a between subjects factor in the statistical design. The threshold for significance was p < 0.05.

## Results

**Acute administration** of 20 mg/kg pentilludin in PEG600 to rats (n = 3/timepoint) provided significant plasma levels following iv or po administration (Table I). There was evidence for a modest terminal half-life of plasma pentilludin between one and two hours. Data from 0.5 to 4h support 15% bioavailability for po pentilludin in PEG600. Rats treated acutely with 20 mg/kg pentilludin displayed no signs of gross toxicity.

**Table I.**
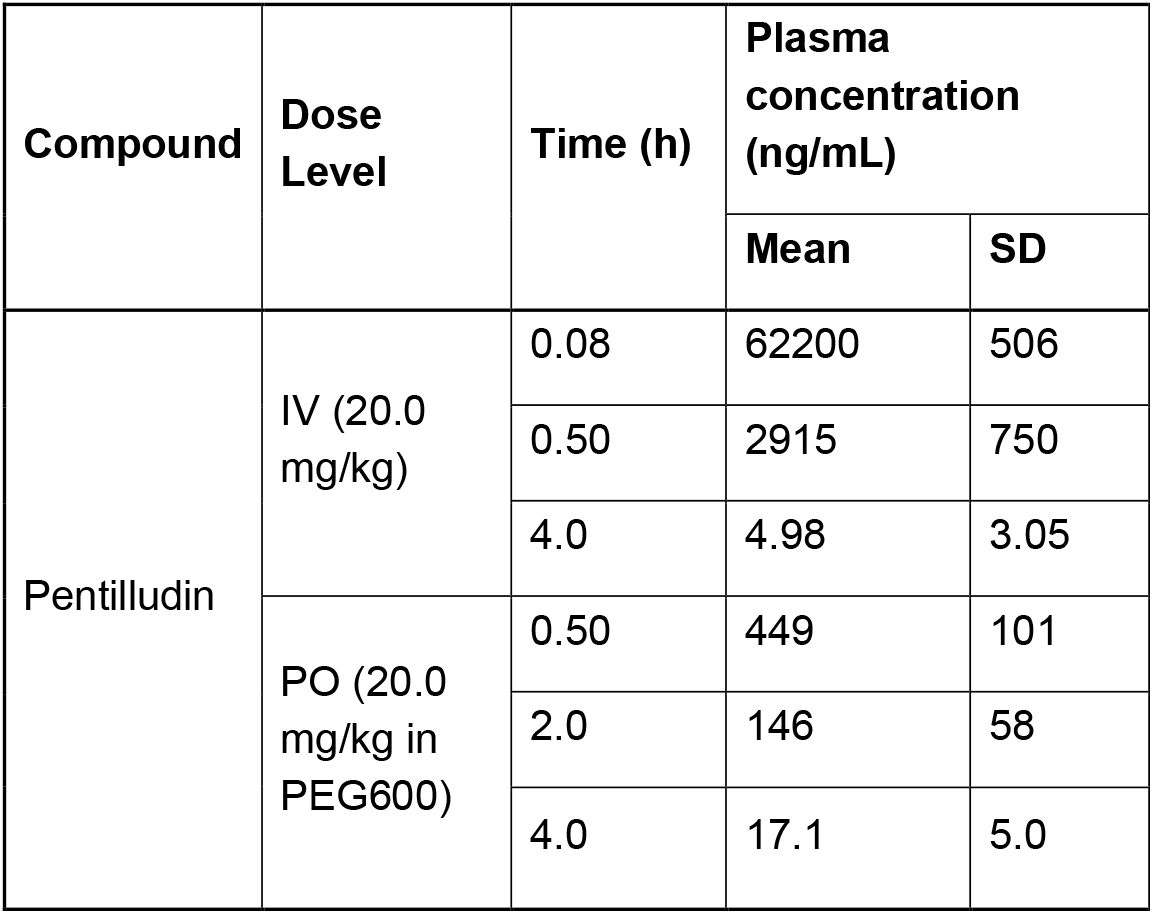
Plasma pentilludin levels following iv or po administration of 20 mg/kg in PEG600.

### Repeated two week daily administration

Two-week daily oral dosing of pentilludin at 50 or 75 mg/kg in PEG600 produced no significant effects on the observational battery, weight, plasma chemistries or hematological assessments (Fig 2).

**Fig 2.**
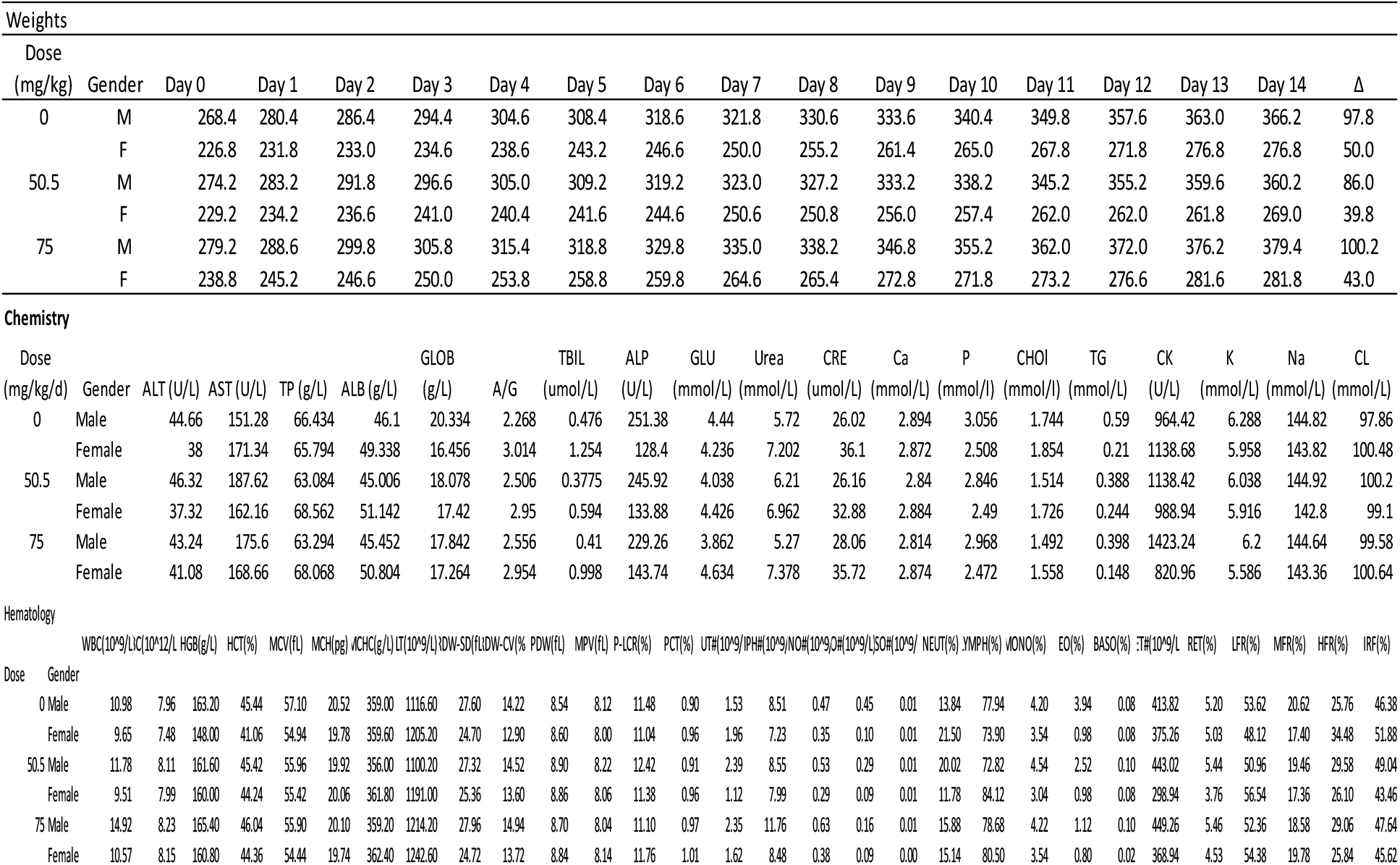
Weights (daily) and serum chemistries and hematology (values on day 14) for rats treated with PEG600 vehicle or pentilludin doses (50 and 75 mg/kg) po for two weeks

There were no pentilludin-associated effects detected on histopathology in our good laboratory practices toxicology study. Several of the rats treated vehicle and several treated with 50 or 75 mg/kg/d po doses for two weeks basophilic kidney histopathology found in more extensive form in chronic progressive nephropathy (CPN). Such idiopathic basophilic kidney lesions have been identified in 1) aging rats and 2) rats treated with a number of currently licensed drugs that do not produce these abnormalities in dogs or humans (20).

In more recent studies of rats treated for two weeks with 100 and 150 mg/kg/d po, there was proteinuria in rats treated with 150 mg/kg/d that recovered within two weeks of the end of pentilludin administration.

Otherwise, our good laboratory practices work replicated the lack of effect of pentilludin on plasma chemistry, hematology and behavioral observation batteries noted in nonGLP studies.

### Self-administration

Effects of pentilludin on self-administration of the stimulant amphetamine and the short acting opiate agonist remifentanil were then tested in rats. Due to the irreversible mechanism of action of pentilludin, we administered pentilludin or vehicle prior to the start of alternating self-administration sessions 2-3 days per week.

#### Amphetamine

Administration of pentilludin sc to rats that have been trained to self-administer amphetamine *iv* prior to every-other self-administration session robustly reduced this amphetamine self-administration (Fig 3). As anticipated, pentilludin administration prior to one session (“treatment”, Fig 3) continues to reduce self-administration during the next session 2-3 days later (“post”), consistent with the irreversible mechanism of action noted *in vitro* (4). We thus pretreated with pentilludin or vehicle prior to every other self-administration session, providing data from 6 pairs of sessions.

**Fig 3.**
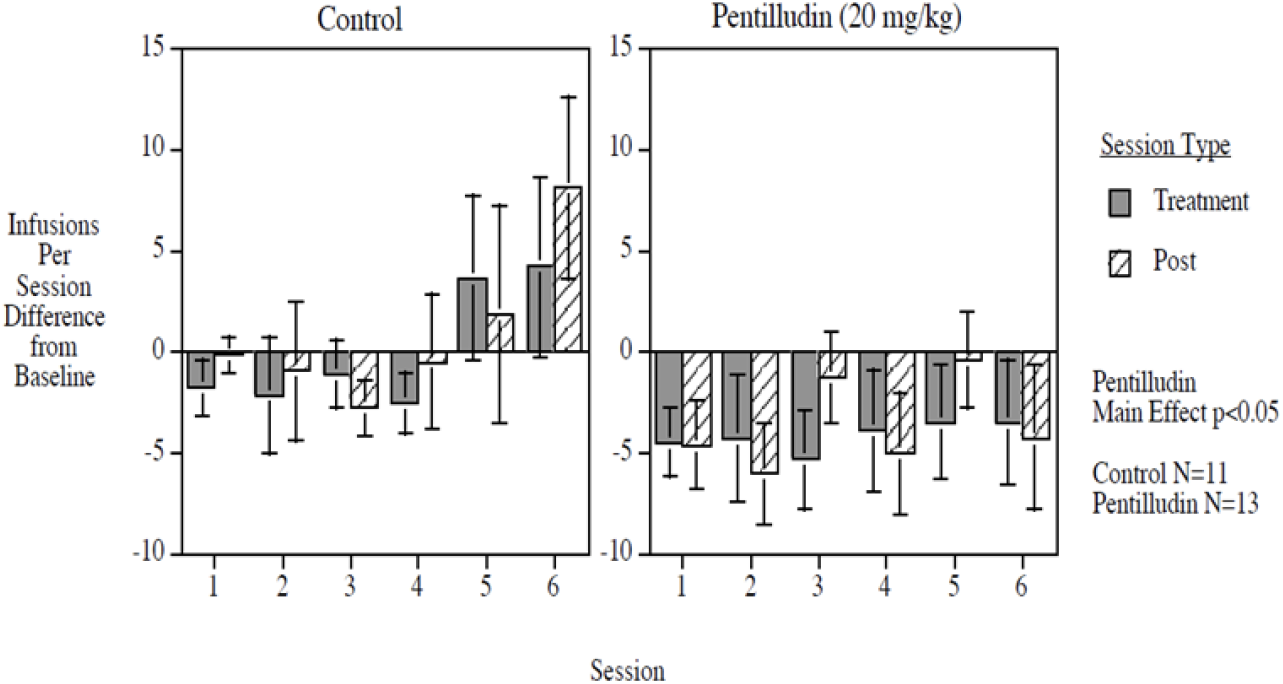
Effects of pentilludin or vehicle pretreatment (20 mg/kg prior to odd numbered sessions) on amphetamine seif administration (values are difference from baseline)

Pentilludin provided a significant main effect, reducing self-administration by 6.2 infusions per session overall relative to controls (F (1,22) = 4.42, p< 0.05). It virtually eliminated the escalation of amphetamine self-administration noted during the course of the 12 sessions in the vehicle-pretreated rats. During the last two of the total 12 post-acquisition self-administration sessions, rats pretreated with vehicle before the penultimate session self-administered at 87% increase from their last acquisition sessions. By contrast, rats pretreated with pentilludin sc self-administered at a 23% decrease from their last acquisition sessions.

There was no evidence for motor or other disabilities that would confound these results. Rates of responding on the control lever that led to no drug being delivered were similar in treated and untreated rats. There were no chronic pentilludin-induced decreases in lever pressing for self-administration of food pellets over eight sessions of testing (control = 77.4 ± 8.6 food pellets, pentilludin = 85.6 ± 6.3). perone-hour session).

#### Remifentanil

We administered pentilludin sc prior to every other self-administration session in rats that had been trained to self-administer remifentanil iv (Fig 4). This alternating session pretreatment significantly attenuated the increase in remifentanil self-administration over the course of 12 self-administration sessions (6 pairs of drug and post drug sessions). There was a significant pentilludin x session block interaction (F (5,90) = 3.93, p < 0.005; exact p = 0.0029). The pentilludin-treated rats had a significantly lower slope of increasing remifentanil self-administration across session blocks compared with vehicle-pretreated rats (F1,18) = 6.09, p < 0.025; exact p=0.024). By the end of testing during the 11^th^ and 12^th^ sessions, the pentilludin-pretreated rats displayed 26.5% smaller increase in remifentanil self-administration relative to vehicle-treated controls. Responding on the control lever did not differ in pentilludin *vs* vehicle-treated rats.

**Fig 4.**
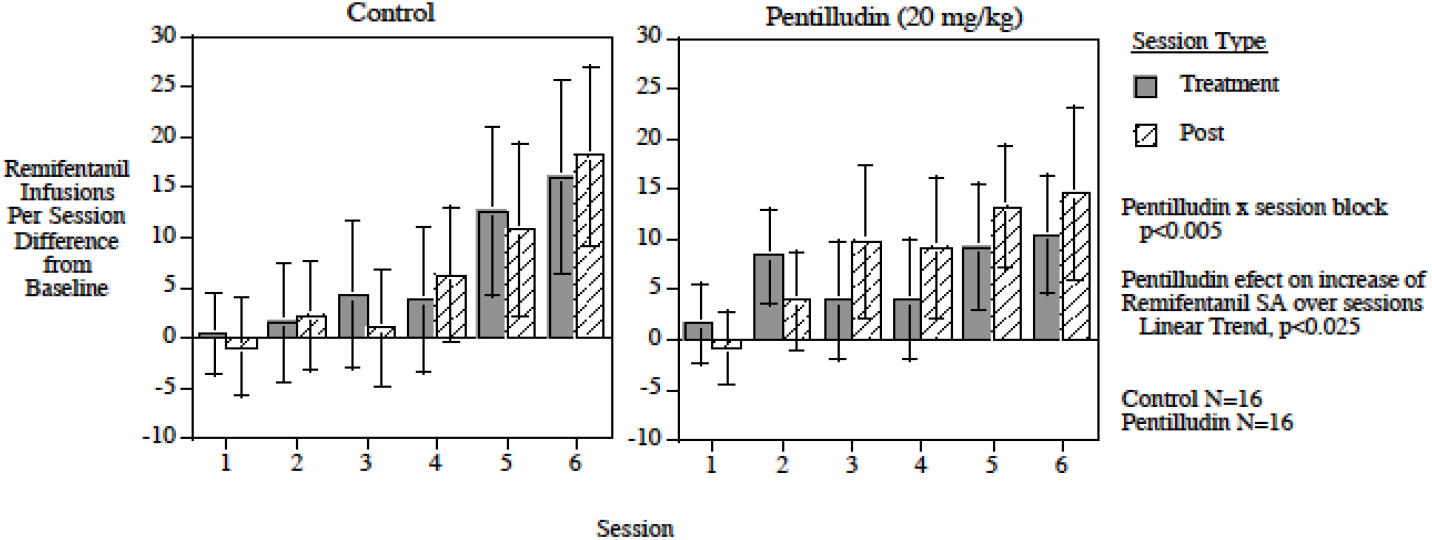
Effects of pentilludin or vehicle pretreatment (20 mg/kg prior to odd numbered sessions) on remifentanil seif administration (values are difference from baseline)

## Discussion

Our results with pentilludin validate use of rats to study the effects of this potent PTPRD phosphatase inhibitor. These data document pentilludin efficacy in reducing intravenous self-administration, a measure of reward from a stimulant and (modestly) an opiate, in this species.

Our results substantially extend previously reported toxicologic and pharmacologic studies in mice (1, 21). Mice tolerate weeks-long repeated treatments with our lead PTPRD phosphatase inhibitor, 7-BIA or with pentilludin with little evidence for toxicity at pentilludin doses up to 100 mg/kg/d po. Treatment with 7-BIA also reduces stimulant reward in mice, as assessed by both cocaine conditioned place preference and cocaine self-administration (4).

Our results also complement and extend human and mouse genetic studies. As noted above, we initially identified and others have confirmed PTPRD as a target for antiaddiction therapeutic development based on modest but repeated genetic associations between variants at the PTPRD locus and human individual differences in 1) vulnerability to develop a substance use disorder (polysubstance use (7-9), opioid use disorder (10) and alcohol use disorder (11)); 2) ability to quit smoking (12, 13); 3) ability to quit use of opioids (14); and 4) ability to reduce alcohol use (when aided by naltrexone) (15). PTPRD SNPs display 10^−6^ < p < 10^−7^ association with a specific constellation of rewarding responses to amphetamine administered po in a human laboratory setting (16)(22).

Heterozygous PTPRD knockout mice with 50% of wildtype levels of PTPRD expression display substantially reduced reward from stimulants (4, 5). Genetic variants at the PTPRD locus are associated with about 60% individual differences in the levels of PTPRD expression in postmortem brains (5), helping to validate the relevance of these mouse results in light of this human data.

Illudalic acid analogs including pentilludin and 7-BIA are thought to irreversibly inhibit PTPRD’s phosphatase (23), supporting a model in which the physiological half-lives of their actions are likely determined by the half-life of PTPRD in the plasma membrane rather than the pharmacological half life of pentilludin or 7-BIA in plasma. Our data support good efficacy of pentilludin dosing prior to odd-numbered sessions to reduce self-administration during these sessions and during the next (even numbered) sessions 2-3 days later. These results fit well with the 1.7-day half-life estimates reported for PTPRD in neuronal cultures (24). Twice-weekly dosing, a benefit of irreversible action, might have advantages in potential therapeutic settings.

We found basophilic lesions in the kidneys of a number of the rats treated with daily doses of vehicle and with pentilludin for two weeks, as we had found in mice (1). None of the rats treated with doses up to 100 mg/kg/d displayed alterations in serum chemistries related to renal function. Similar lesions, reminiscent of chronic progressive nephropathy, are found in older rats and mice and in rats and mice treated with a number of drugs that are well tolerated by humans (20). In our further experiments, rats treated with 150 mg/kg/d did develop modest and reversible proteinuria, again with no serum chemistry abnormalities in measures relating to renal function. Further, dogs treated with the equivalent dose (15 mg/kg/d po) for two weeks fail to display any kidney pathology. Failure of these chronic progressive nephropathy-like lesions to alter serum chemistries or to be found in dogs treated with equivalent doses suggests that this intermittent pathology does not invalidate rats as appropriate models for study of pentilludin effects on reward from stimulants or opiates.

Results from pentilludin effects in rat and human test systems support use of rat data to propose effects of this drug that might be found in humans. There is only modest metabolism when pentilludin is exposed to liver microsomes from either rats or human. By contrast, there is robust metabolism when pentilludin is exposed to plasma from each of these species. In our initial work, recombinant paraoxonase interacts avidly with pentilludin, providing a likely mechanism for this plasma degradation of pentilludin. These similar metabolic patterns also validate use of rat data to foreshadow likely pentilludin effects in humans.

We have found larger effects of pentilludin pretreatment on amphetamine self-administration than on remifentanil self-administration. This difference has motivated us to target development of pentilludin to aid abstinence by individuals seeking treatment for stimulant use disorders (3). Since opiate use is common among individuals with stimulant use disorders, our data that shows no increase in opiate self-administration after pentilludin treatment is comforting. Conceivably, pentilludin effects on opiate reward in humans might even exceed the modest reductions in rat opiate self-administration that we note in the current work. Such larger effects might receive support from the relatively robust reported genetic associations of human PTPRD variants with individual differences in ability to reduce use of opiates (10, 14).

In summary, the present work validates rats as a species in which to test pentilludin effects. It documents sizable pentilludin effects on amphetamine seif administration as well as more modest effects on remifentanil self-administration. Our results support a timecourse of pentilludin physiological action that exceeds the timecourse predicted by its plasma level pharmacologic activities, as anticipated if pentilludin irreversibly inhibited the actions of a plasma membrane protein with a plasma membrane half live of between 1-2 days. Our results support continued development of pentilludin to reduce reward from stimulants in humans.

## Acknowledgements

We are grateful for substantial assistance and advice from Drs Jane Acri, Matt Seager, Nate Appel, Rik Klein, Jia-bei Wang and Dan Deaver, to contributions of students Asha Barnes, Cindy Su, Emoni Barbour, Jose Jarquin, Kennedy Hill, Maggie Dercole, Rongxi Fan, Wyatt Pfau, Megan Stout and Noe Jr Bravo Velazquez during the course of this work and to contributions of Jerry Wang and colleagues (Netchem), Nouara Sadaoui, Jingjing Zhang, Wanyong Feng and their colleagues (Bioiduro-Sundia) and Sam Hendrix and colleagues (CARE/Mountain West Research).

## Author Contributions

GRU: Substantial contributions to the conception and design of the work, analysis and interpretation of data for the work, drafting the work and revising it critically for important intellectual content; final approval of the version to be published, agreement to be accountable for all aspects of the work in ensuring that questions related to the accuracy or integrity of any part of the work are appropriately investigated and resolved.

BK, JC, EL: Substantial contributions to the acquisition, analysis and interpretation of data for the work, revising it critically for important intellectual content.

IH: Substantial contributions to management of contractual work, analysis and interpretation of data for the work, revising it critically for important intellectual content.

BG, JS, CW: Substantial contributions to acquisition of data

## Funding

We are grateful for support from NIH/NIDA UO1DA047713 and UG3DA056039 for this work.

## Competing Interests

Pentilludin is covered by US patent 11,987,564 issued to the US Department of Veterans Affairs (G Uhl, I Henderson, W Wang and T Prisinzano, inventors).

No other competing interests: G Uhl, B Kannan, J Choi

EL receives funding for nicotine research from Philip Morris International and serves as consultant to Proctor and Gamble for developmental neurotoxicology in zebrafish.

## Notes

### Competing Interest Statement

George Uhl is a coinventor of VA intellectual property that covers pentilludin

## References

1. Henderson IM, Zeng F, Bhuiyan NH, Luo D, Martinez M, Smoake J, et al. Structure-activity studies of PTPRD phosphatase inhibitors identify a 7-cyclopentymethoxy illudalic acid analog candidate for development. Biochem Pharmacol. 2022;195:114868.

2. Henderson IM, Marez C, Dokladny K, Smoake J, Martinez M, Johnson D, et al. Substrate-selective positive allosteric modulation of PTPRD’s phosphatase by flavonols. Biochem Pharmacol. 2022;202:115109.

3. Uhl GR. Selecting the appropriate hurdles and endpoints for pentilludin, a novel antiaddiction pharmacotherapeutic targeting the receptor type protein tyrosine phosphatase D. Front Psychiatry. 2023;14:1031283.

4. Uhl GR, Martinez MJ, Paik P, Sulima A, Bi GH, Iyer MR, et al. Cocaine reward is reduced by decreased expression of receptor-type protein tyrosine phosphatase D (PTPRD) and by a novel PTPRD antagonist. Proc Natl Acad Sci U S A. 2018;115(45):11597–602.

5. Drgonova J, Walther D, Wang KJ, Hartstein GL, Lochte B, Troncoso J, et al. Mouse model for PTPRD associations with WED/RLS and addiction: reduced expression alters locomotion, sleep behaviors and cocaine-conditioned place preference. Mol Med. 2015.

6. Uhl GR, Drgonova J. Cell adhesion molecules: druggable targets for modulating the connectome and brain disorders? Neuropsychopharmacology. 2014;39(1):235.

7. Liu QR, Drgon T, Walther D, Johnson C, Poleskaya O, Hess J, et al. Pooled association genome scanning: validation and use to identify addiction vulnerability loci in two samples. Proc Natl Acad Sci U S A. 2005;102(33):11864–9.

8. Liu QR, Drgon T, Johnson C, Walther D, Hess J, Uhl GR. Addiction molecular genetics: 639,401 SNP whole genome association identifies many “cell adhesion” genes. Am J Med Genet B Neuropsychiatr Genet. 2006;141B(8):918–25.

9. Drgon T, Johnson CA, Nino M, Drgonova J, Walther DM, Uhl GR. “Replicated” genome wide association for dependence on illegal substances: genomic regions identified by overlapping clusters of nominally positive SNPs. Am J Med Genet B Neuropsychiatr Genet. 2011;156(2):125–38.

10. Li D, Zhao H, Kranzler HR, Li MD, Jensen KP, Zayats T, et al. Genome-wide association study of copy number variations (CNVs) with opioid dependence. Neuropsychopharmacology. 2015;40(4):1016–26.

11. Jung j ZH, Grant BF, Chou P Identification of novel genetic variants of DMS5 alcohol use disorder: Genome wide association study in National Epidemiological Survey on Alcohol Related Conditions-III. Abstracts, World Congress of Psychaitroc Genetics. 2017.

12. Uhl GR, Liu QR, Drgon T, Johnson C, Walther D, Rose JE. Molecular genetics of nicotine dependence and abstinence: whole genome association using 520,000 SNPs. BMC Genet. 2007;8:10.

13. Uhl GR, Liu QR, Drgon T, Johnson C, Walther D, Rose JE, et al. Molecular genetics of successful smoking cessation: convergent genome-wide association study results. Arch Gen Psychiatry. 2008;65(6):683–93.

14. Cox JW, Sherva RM, Lunetta KL, Johnson EC, Martin NG, Degenhardt L, et al. Genome-Wide Association Study of Opioid Cessation. J Clin Med. 2020;9(1).

15. Biernacka JM, Coombes BJ, Batzler A, Ho AM, Geske JR, Frank J, et al. Genetic contributions to alcohol use disorder treatment outcomes: a genome-wide pharmacogenomics study. Neuropsychopharmacology. 2021;46(12):2132–9.

16. Hart AB, Engelhardt BE, Wardle MC, Sokoloff G, Stephens M, de Wit H, et al. Genome-wide association study of d-amphetamine response in healthy volunteers identifies putative associations, including cadherin 13 (CDH13). PLoS One. 2012;7(8):e42646.

17. A Hart and A Palmer, personal communication, 2013

18. Mishra I, Xie WR, Bournat JC, He Y, Wang C, Silva ES, et al. Protein tyrosine phosphatase receptor delta serves as the orexigenic asprosin receptor. Cell Metab. 2022;34(4):549–63 e8.

19. O’Connor EC, Chapman K, Butler P, Mead AN. The predictive validity of the rat self-administration model for abuse liability. Neurosci Biobehav Rev. 2011;35(3):912–38.

20. Hard GC, Johnson KJ, Cohen SM. A comparison of rat chronic progressive nephropathy with human renal disease-implications for human risk assessment. Crit Rev Toxicol. 2009;39(4):332–46.

21. Uhl GR, Martinez MJ. PTPRD: neurobiology, genetics, and initial pharmacology of a pleiotropic contributor to brain phenotypes. Ann N Y Acad Sci. 2019;1451(1):112–29.

22. A Hart and A Palmer, personal communication, 2013

23. Ling Q, Huang Y, Zhou Y, Cai Z, Xiong B, Zhang Y, et al. Illudalic acid as a potential LAR inhibitor: synthesis, SAR, and preliminary studies on the mechanism of action. Bioorg Med Chem. 2008;16(15):7399–409.

24. Heo S, Diering GH, Na CH, Nirujogi RS, Bachman JL, Pandey A, et al. Identification of long-lived synaptic proteins by proteomic analysis of synaptosome protein turnover. Proc Natl Acad Sci U S A. 2018;115(16):E3827–E36.

